# Infection by the lungworm *Rhabdias pseudosphaerocephala* affects the expression of immune-related microRNAs by its co-evolved host, the cane toad *Rhinella marina*

**DOI:** 10.1101/2023.06.19.545515

**Authors:** Tsering C. L. Chan, Boris Yagound, Gregory P. Brown, Harrison J. F. Eyck, Richard Shine, Lee A. Rollins

## Abstract

Parasites may suppress the immune function of an infected host using microRNAs (miRNAs) to prevent protein production. Nonetheless, little is known about the diversity of miRNAs and their mode(s) of action. In this study, we investigated the effects of infection by a parasitic lungworm (*Rhabdias pseudosphaerocephala*) on miRNA and mRNA expression of its host, the invasive cane toad (*Rhinella marina*). We compared miRNA and mRNA expression in naïve toads that had never been infected by lungworms to toads that were infected with lungworms for the first time in their lives, and to toads that were infected the second time in their lives (i.e., had two consecutive infections). In total, we identified 434 known miRNAs and 106 potential novel miRNAs. Compared to uninfected toads, infected animals upregulated five (single-infection treatment) or four (multiple-infection treatment) miRNAs. Seven of these differentially expressed miRNAs were associated with gene pathways related to the immune response, potentially reflecting immunosuppression of cane toads by their parasites. Infected hosts did not respond with substantial mRNA transcription, with only one differentially expressed gene between control and single-infection hosts. Our study suggests that miRNA-mediated interactions may play a role in mediating the interaction between the parasite and its host. Our findings clarify the role of miRNAs in host-parasite interactions, in a system in which an ongoing range expansion by the host has generated substantial divergence in host-parasite interactions.

## Introduction

Host-parasite interactions are ubiquitous in nature and typically result from complex coevolutionary dynamics (Anderson & May, 1982; D. Ebert & Fields, 2020). As a result, parasites often display adaptations increasing their ability to infect, survive and reproduce within their hosts through directly manipulating host behaviour and/or immune function (Coakley, Buck, & Maizels, 2016; Maizels, 2020; Rambani et al., 2020). The immunoregulatory effects of parasites can persist after infection has been lost, reducing the reactivity of the host’s immune system to non-parasite stimuli such as allergens (Johnston, McSorley, Anderton, Wigmore, & Maizels, 2014; Rodrigues et al., 2008).

The ability of a parasite to successfully complete its life cycle often relies on molecular mechanisms, such as microRNAs (miRNA), that can directly manipulate host gene function (Ghalehnoei, Bagheri, Fakhar, & Mishan, 2020; Han et al., 2013; He & Pan, 2022; Rambani et al., 2020; Simo, Lueong, Grebaut, Guny, & Hoheisel, 2015). miRNAs are non-coding RNAs that regulate gene expression by binding to a target mRNA (Campbell et al., 2015). This results in the suppression of protein production by repressing translation or degrading target mRNA (Baek et al., 2008). miRNAs have been found in all animal model systems and are critical for regulating genes in a variety of biological processes (O’Brien, Hayder, Zayed, & Peng, 2018), including innate and adaptive immune responses (Carissimi, Fulci, & Macino, 2009). miRNAs can also promote stability in gene expression by buffering the effects of fluctuations in mRNA concentration (M. S. Ebert & Sharp, 2012). A single miRNA can potentially mediate many genes, and multiple miRNAs may regulate the same target gene, enabling them to dynamically affect downstream biological functions (Selbach et al., 2008). miRNA sequences are highly conserved across species (Ghosh, Chakrabarti, & Mallick, 2007). This facilitates cross-species interactions between miRNAs and miRNA-mediated functions, such as the release of miRNAs into the host system (Buck et al., 2014; He & Pan, 2022; Weinstock & Elliott, 2013). For example, the parasitic apicomplexan *Cryptosporidium parvum* utilises host miRNA pathways to induce the death of activated T cells and boost parasite survival despite the host’s immune response (Zheng, Cai, & Bradley, 2013). Conversely, the host defence system can reduce parasite survival through targeted manipulation of parasite miRNAs, disrupting key biological functions (Maizels, 2020). LaMonte *et al*. (2012) observed that miRNAs found in human red blood cells affected by sickle cell disease contribute to resistance against the malaria parasite *Plasmodium falciparum* by reducing parasite growth and infection rate. Additionally, *Plasmodium* parasites were found to have incorporated host miRNA into their mRNA transcripts, inhibiting parasite mRNA translation. These findings illustrate the complex two-way role of miRNAs in host-parasite interactions.

Helminths (i.e., nematodes and platyhelminths) are arguably the best-studied examples of parasitic metazoans that depend on molecular manipulation of their hosts. During infection, helminths not only release proteins that bind to host cellular receptors, but they also release exosomes (i.e., extracellular vesicles) containing miRNAs that can alter host gene expression (Coakley et al., 2016). Helminths often disrupt multiple pathways involving many genes to suppress host immune responses (Entwistle & Wilson, 2017; Johnston et al., 2014; Raissi et al., 2020; Rambani et al., 2020; Simo et al., 2015). Conversely, hosts can limit the damage caused by infection through the release of their own regulatory miRNAs that negatively affect helminth parasites (Yuan, Luo, Xu, Zeng, & Wu, 2019).

The invasive cane toad (*Rhinella marina*) is a prime candidate for investigating parasitic manipulation of host miRNA and mRNA. Cane toads were introduced to Australia in 1935, following successive translocations from South America (Easteal, 1981). This translocation history allowed toads to escape many of the parasites found in their native range (Campiao et al., 2014; Selechnik, Rollins, Brown, Kelehear, & Shine, 2017). Of the helminths that parasitise native cane toads, only one nematode species, the lungworm *Rhabdias pseudosphaerocephala*, is thought to have persisted through these introductions (Dubey & Shine, 2008; Kuzmin, Tkach, & Brooks, 2007). *Rhabdias pseudosphaerocephala* is the most common lung parasite in Australian cane toads (Phillips et al., 2010), and is much less effective at infecting native Australian frogs (Pizzatto, Shilton, & Shine, 2010). Conversely, native Australian *Rhabdias* species are also ineffective against invasive cane toads, which indicates that *R. pseudosphaerocephala*’s ability to overcome the cane toad’s immune defences results from a long coevolutionary history (F. B. Nelson, Brown, Shilton, & Shine, 2015a; F. B. L. Nelson, Brown, Dubey, & Shine, 2015b).

*Rhabdias pseudosphaerocephala* has no intermediate host species, and adult lungworms remain in the same individual toad throughout their lives. Adult parasites are attached to the host’s lungs and lay eggs, which are coughed into the throat by the host and swallowed into the digestive system (Crystal Kelehear, Webb, Hagman, & Shine, 2011a). The eggs hatch into L1 larvae in the colon, and are excreted by the host, whereupon they live freely in the soil as sexually reproductive adults (Baker, 1979). After mating, their offspring (L3 larvae) develop for approximately four to seven days inside the female L1 larva, before breaking open the maternal cuticle, releasing themselves into the environment (Baker, 1979). L3 larvae infect cane toads by entering through the skin or eye socket, after which they must travel to the lungs. In this phase, they are exposed to the blood plasma of the host (Brown, Phillips et al. 2015). The plasma contains host immune products and reduces the longevity of L3 larvae (Mayer, Shine et al. 2021a). Once inside the lungs, larvae feed on epithelium and mature into adults in five or more days (C. Kelehear, Brown, & Shine, 2012). L3 larvae are capable of infecting metamorph, juvenile and adult toads (C. Kelehear, Brown, & Shine, 2011b; C. Kelehear, Webb, & Shine, 2009a). Toads may also become infected by consuming infected prey or L3 larvae (C. Kelehear et al., 2009a).

The molecular mechanisms at play in an infection by *R. pseudosphaerocephala* remain uncertain. On the one hand, these lungworms can negatively impact the fitness of their hosts. Indeed, high parasite loads can cause declines in the growth rates, boldness, cardiac capacity, and activity levels of toads (Finnerty, Shilton, Shine, & Brown, 2017; C. Kelehear et al., 2011b; Selechnik et al., 2017). Infection may also cause physical damage from entry through the skin or the migration of L3 larvae through tissue once inside the host (C. Kelehear et al., 2011b). On the other hand, a specific immune response to *R. pseudosphaerocephala* infection has not been observed in cane toads, suggesting that this parasite is tolerated by their hosts (Crystal Kelehear et al., 2011a). Dead L3 larvae have infrequently been found in toad tissues surrounded by granulomas, indicating an activation of the host’ immune system, but it is unclear whether this defence response is responsible for the death of the parasitic larvae (Pizzatto et al., 2010). Further, natural levels of lungworm infection as seen in the wild appear to have minor impacts on toad performance and survivability, where wild individuals carrying more than 200 *R. pseudosphaerocephala* have been observed (F. B. L. Nelson et al., 2015b). Chronic stimulation of the immune response can negatively affect metabolic rate and fecundity (Bashir-Tanoli & Tinsley, 2014). In the wild, there is a high chance of a cane toad becoming infected by *R. pseudosphaerocephala*, because prevalence of this parasite is 70– 80% in established toad populations, although populations on the edge of the expanding range are relatively parasite-free (Lettoof, Greenlees, Stockwell, & Shine, 2013). It is possible, given the cost of mounting a repeated immune response and the high likelihood of encountering *R. pseudosphaerocephala* in most cane toad populations, that selection for resistance to this parasite is limited (Phillips et al., 2010).

Cane toads also employ anti-parasite behaviours to avoid infection by these macro-parasites, because they have been observed attempting to dislodge L3 larvae by kicking, flicking their tongues, and blinking although these methods are largely ineffective in preventing infection (C. Kelehear et al., 2011b; Crystal Kelehear et al., 2011a). The success of *R. pseudosphaerocephala* implies they are able to evade the host immune system, either through immunosuppression or by masking themselves, such as through coating themselves in mucus found in the lungs to cover their non-host antigens. It is possible that miRNA-mediated immunosuppression from *R. pseudosphaerocephala* during infection contributes to the high prevalence and infection success of this invasive nematode. Nonetheless, this hypothesis has never been investigated. In this study, we aimed to: i) characterise known and novel miRNA profiles of cane toads, ii) investigate changes to miRNA and mRNA expression in cane toads infected with *R. pseudosphaerocephala* to examine whether infection triggers miRNA-mediated innate or adaptive immune response pathways, iii) determine whether different exposure regimes (i.e., naïve vs non-naïve hosts) affect miRNA and mRNA expression, and iv) determine whether there is evidence of parasite miRNAs within host tissues.

## Materials and Methods

### Sample collection and infection treatments

We collected six pairs of adult cane toads from three localities spread across the invasive range in Australia: two pairs from Queensland (17.00°S, 145.44°E), two pairs from the Northern Territory (12.58°S, 131.32°E), and two pairs from Western Australia (14.79°S 125.83°E; Table S1). We induced G0 toads to breed by injection of leuprorelin acetate (Lucrin; Abbott Australasia, Australia). We collected toad eggs and raised tadpoles to metamorphosis in a common garden setting in our field station in Middle Point (Northern Territory; see DeVore et al. 2021 for detailed methods).

We allocated 24 G1 female toads from across pairs from the three collection localities to one of three treatment groups (Figure 1A): i) no infection (i.e., control response), ii) single lungworms infection episode (i.e., naïve host response, to investigate innate immune responses), or iii) two consecutive lungworm infection episodes (i.e., non-naïve host response, to investigate both innate and acquired immune responses; Table S1). To obtain infectious L3 larvae, we collected adult *R. pseudosphaerocephala* from the lungs of infected Northern Territory toads. We harvested lungworm eggs and cultured them in a mixture of water and toad faeces until the larvae reached the infectious L3 stage. We exposed single-infected toads to 30 L3 larvae only once at 19 months of age. We exposed double-infected toads twice: once to 50–100 L3 larvae at nine months of age, then again to 30 L3 larvae 10 months later when the initial infections had naturally cleared (verified by lack of lungworm larvae passing in faeces).

**Figure 1.**
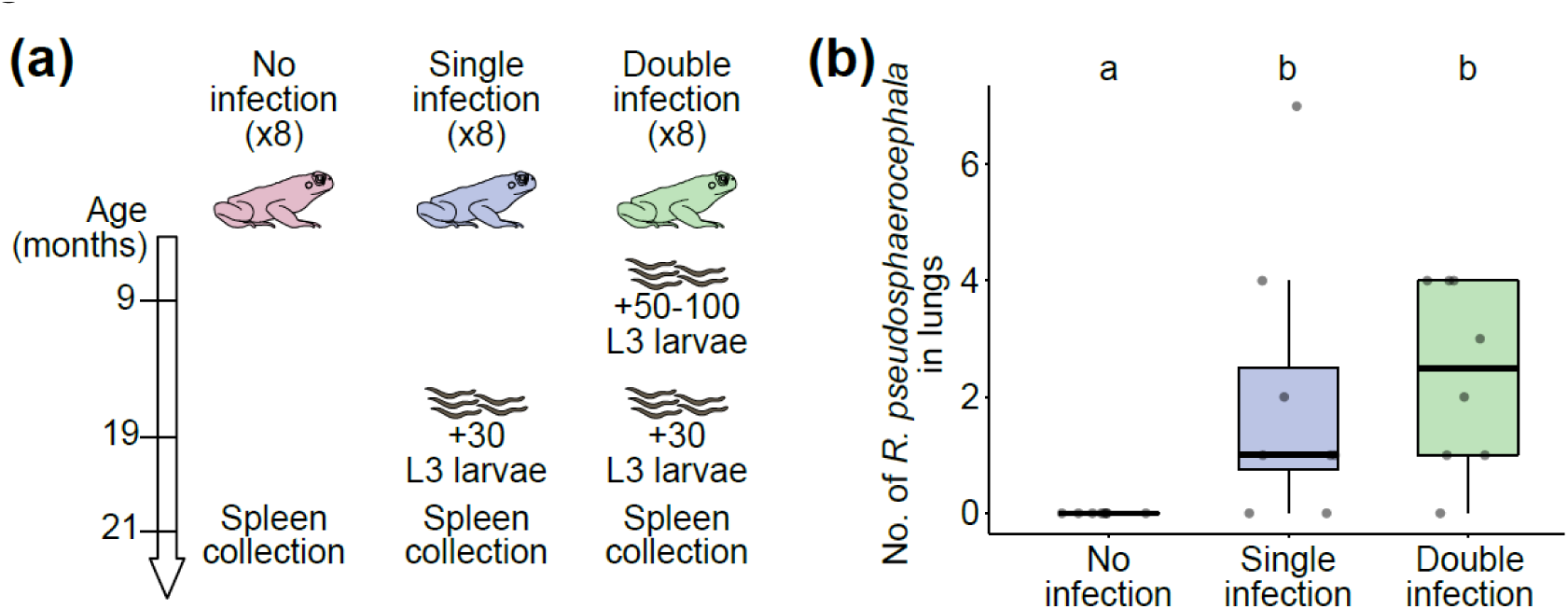
Study design. (**a**) Female cane toads were allocated to one of three treatment groups (8 individuals each): i) no infection (i.e., control response), ii) single infection with 30 L3 larvae at 19 months (i.e., naïve host response), or iii) double infection, once with 50–100 L3 larvae at 9 months, then again with 30 L3 larvae at 19 months (i.e., non-naïve host response). Spleens were collected for RNA extraction at 21 months for all individuals. (**b**) Number of *R. pseudosphaerocephala* parasites found in lungs in each treatment groups. Different letters denote statistical differences (Kruskal-Wallis rank sum test followed by Dunn’s post hoc tests).

To expose toads to infectious larvae, we placed toads individually into clean 500 mL containers lined with filter paper. We added L3 larvae in 300 µL of water to the container with the toad and left them for 18 h. We then removed toads from the infection chambers and housed them individually in 15 L containers for the duration of the experiment. We also held control toads in infection chambers for 18 h, but without introducing any L3 larvae.

Two months after the last possible infection period (i.e., when toads were 21 months old), we euthanized them with an injection of sodium pentobarbital and then dissected them to count the number of adult nematodes in their lungs. We collected spleens immediately after euthanasia, preserved them in RNALater (Qiagen, USA), and stored them at –80°C. We chose spleens for RNA extraction because this organ is involved in host infection response and filters blood (Mebius & Kraal, 2005), where circulating parasite miRNAs are likely to be present.

### RNA extraction and sequencing

We disrupted whole spleens by crushing them with mortar and pestle using liquid nitrogen, then homogenised them in QIAzol lysis reagent (Qiagen) by passing them through a sterile 20-gauge needle. The amount of tissue used varied between individuals (<1 mg to 35 mg). We extracted total RNA using the miRNeasy Mini Kit (Qiagen) following manufacturer’s instructions. We quantified RNA concentrations and quality using a Qubit 4 Fluorometer (Thermo Fisher Scientific) using the Qubit RNA High Sensitivity (HS) kit (Thermo Fisher Scientific) and Qubit RNA IQ Assay kit (Thermo Fisher Scientific). We submitted samples to the Ramaciotti Centre for Genomics (University of New South Wales, Sydney, Australia) for miRNA library construction and sequencing. Average RNA Integrity Number values for all samples were 8.9 ± 0.4 (mean ± SD; Table S1). For miRNA library construction, we used 100 ng of extracted miRNA for the QIAseq miRNA UDI Library protocol per manufacturer’s instructions. We then performed 75 bp single-end sequencing using an Illumina NextSeq 500 system. For mRNA library construction, we used the Illumina Stranded mRNA Prep, Ligation kit (Illumina), followed by 100 bp single-end sequencing on an Illumina NovaSeq 6000 SP system.

### Data processing

We checked miRNA reads quality with FastQC 0.11.8 (Andrews, 2010).We used Cutadapt 2.10 (Martin, 2011) to remove adapters, reads with length <18 and >24 nt, and low-quality reads (Phred score <30). We aligned clean reads using Bowtie 1.2.0 (Langmead, 2010) with the –v alignment mode (settings: 1 mismatches allowed, using –best and –strata reporting modes, as recommended by Ziemann, Kaspi, and El-Osta (2016) for mapping short reads) against all available mature miRNA sequences from miRbase 22.1 (Griffiths-Jones, Grocock, Van Dongen, Bateman, & Enright, 2006). We grouped miRNAs in miRNA families following Li et al. (2010), as members of the same miRNA family usually have similar physiological functions despite differing in structure (Griffiths-Jones, Bateman, Marshall, Khanna, & Eddy, 2003).

To detect novel miRNAs, we mapped reads that did not match to known miRNAs within the database against other small RNA databases using Bowtie (settings: 0 mismatches allowed, using –best and –strata reporting modes) –high confidence tRNA sequences (*Xenopus tropicalis* 9.1, GtRNAdb), ribosomal RNA sequences (SILVA 138.1), and repetitive DNA (RepBase 27.03)– to filter out non-miRNA sequences. We then aligned the remaining reads using mirDeep2 2.0.1.2 (Friedlander, Mackowiak, Li, Chen, & Rajewsky, 2012) through the Galaxy bioinformatics workflow to discover putative novel miRNAs, using default settings. We used the cane toad genome 2.2 (Edwards et al., 2018) and mature *X. tropicalis* miRNAs (miRbase 22.1) as the input databases. We filtered unannotated miRNA sequences for a miRDeep score ≥ 10, a read depth ≥ 10, a significant RNA fold (i.e., estimated probability that the miRNA candidate is a true positive) ≥ 90%, and no Rfam small RNA database alert that could indicate a match with a known non-miRNA sequence. We interpreted sequences which were present in ≥ 10 samples, or ≥ 5 samples within the same treatment group, as potential novel miRNAs. There was a similar number of raw reads, post-trimming reads and mapped reads for control individuals, single-infected individuals and double-infected individuals (Table S2). To compare the presence of miRNA families across different treatments, we discarded families with low read depth (i.e., normalised mean cpm <10).

We quality-checked raw mRNA reads using FastQC. We trimmed adaptor sequences and low-quality reads using Trimmomatic 0.38 (Bolger, Lohse, & Usadel, 2014) with the following parameters: ILLUMINACLIP: path/to/TruSeq3-PE.fa:2:30:10:4 HEADCROP:13 AVGQUAL:30 MINLEN:36 CROP 85. We then mapped trimmed reads to the multi-tissue reference cane toad transcriptome (Richardson et al., 2018) using STAR 2.7.2b (Dobin et al., 2013) in two-pass mode with default parameters. We used Salmon 1.2.1 (Patro, Duggal, Love, Irizarry, & Kingsford, 2017) to quantify gene expression from the resultant BAM files. There was a similar number of raw reads, post-trimming reads and mapped reads for control individuals, single-infected individuals and double-infected individuals (Table S3).

### miRNA differential expression analysis

We kept for differential expression analysis miRNAs with a minimum read count of 10, a minimum total read count across samples of 15, and that appeared in ≥ 50% of samples. We normalised miRNA read counts to trimmed mean of M-values (TMM), as recommended by Tam, Tsao, and McPherson (2015). We used EdgeR 3.9 (Chen, Lun, & Smyth, 2016; Robinson, McCarthy, & Smyth, 2010) for differential expression analysis. To assess the presence of potential outliers we used the limma package (Ritchie et al., 2015) to produce a multi-dimensional scaling plot using normalised and rlog transformed counts, before computing pairwise correlations for all samples (Figure S1). All samples were retained for subsequent analysis.

We conducted pairwise comparisons between all treatment combinations (i.e., control vs single infection, control vs double infection, single infection vs double infection). We adjusted *p*-values for multiple testing (Benjamini & Hochberg, 1995).

### Evidence of parasite miRNA within hosts

We used a twofold approach to determine if there was evidence of parasite miRNA within host tissue. First, we searched mapped sequences for miRNAs that have been described only in nematodes and not vertebrates. Second, we compared known miRNAs that were differentially expressed in infected toads to miRNA infection studies in existing literature determine whether those miRNAs have previously been implicated in parasite manipulation of hosts.

### miRNA target gene prediction

To predict potential gene targets of differentially expressed miRNAs (hereafter, DEmirs), we used the combination of three predicting tools: miRanda (Betel, Koppal, Agius, Sander, & Leslie, 2010), PITA (Kertesz, Iovino, Unnerstall, Gaul, & Segal, 2007) and TargetSpy (Sturm, Hackenberg, Langenberger, & Frishman, 2010), as implemented in sRNAtoolbox (Aparicio-Puerta et al., 2019). We searched target genes in 3’UTR sequences retrieved from the reference cane toad transcriptome (Richardson et al., 2018). We deemed genes as putative targets of a particular miRNA only if predicted so by all three tools (Oliveira et al., 2017). To refine our results, we filtered out putative target genes that had an energy score >–15 kcal/mol (all three tools), and a sequence complementary score <150 (miRanda) (Hitit et al., 2022).

To infer the biological function of putative target genes, we performed Gene Ontology (GO) enrichment analyses using goseq 1.26.0 (Young, Wakefield, Smyth, & Oshlack, 2010). We performed GO analyses using the combined target genes of all DEmirs in each pairwise comparison (i.e., control vs single infection, control vs double infection, single infection vs double infection) using the probability weighting function (PWF) to adjust for transcript length bias, and the Wallenius approximation to test for over-representation. *P*-values were adjusted using the Benjamini-Hochberg method. We used enrichplot (Yu, 2021) to visualise the results.

### mRNA differential expression analysis

We used edgeR to filter out genes with <10 cpm in at least 10 samples. To assess the presence of potential outliers, we normalised and rlog transformed counts, before computing pairwise correlations for all samples. Plotting the resultant correlation matrix with pheatmap 1.0.12 (Kolde, 2019) revealed one potential outlier, toad #476 (Figure S2). To check this result, we carried out differential expression analyses (see below) with and without #476. When #476 was included, we found large differences in gene expression among the three groups (i.e., control vs single infection = 362 differentially expressed genes (hereafter, DEGs); control vs double infection = 0 DEGs; single infection vs double infection: 1,183 DEGs). By contrast, when #476 was excluded, we observed that virtually all significant differences in gene expression levels vanished (with the exception of one DEG between control and single infection, see Results). In other words, it appeared that all the results seen when #476 was included were solely due to that single individual. Until additional data are available, we believe that a cautious approach is warranted before building a narrative around a single cane toad. We thus decided in the main Results section to report the outcome of the differential expression analyses without including toad #476.

We used DESeq2 1.30.1 (Love, Huber, & Anders, 2014) to perform differential expression analyses. Any gene with a Benjamini-Hochberg adjusted *p*-value < 0.05 was deemed significantly differentially expressed.

## Results

### Parasite abundance and infection

As expected, there were no *R. pseudosphaerocephala* parasites in the lungs of any control toads at the time of euthanasia. In contrast, there was a significantly higher parasite load in toads from both infection groups compared to the control group (Kruskal-Wallis rank sum test followed by Dunn’s post hoc tests: both *p* < 0.0073). The number of lungworms in toads from the single and the double infection groups were similar (*p* = 0.25; Figure 1B and Table S1). Individuals in both infection groups that were lacking worms at death (*n* = 3 toads) had similar miRNA expression patterns to those within the same infection treatment group, which suggests that these individuals had been successfully infected (Figure S1). Therefore, these individuals were retained as part of their treatment group on the assumption that the parasites had been cleared from their lungs prior to euthanasia of the host.

### Global and differential miRNA expression

We identified a total of 540 miRNAs, including 434 known miRNAs and 106 potential novel miRNAs (Table S4). The 434 known miRNAs comprised a total of 162 known miRNA families (Figure 2A). Six miRNA families had > 10 members (i.e., let-7, miR-30, miR-29, miR-10, miR-15 and miR-92). Most miRNA families had one or only a few members mapped to miRbase (Figure 2A).

**Figure 2.**
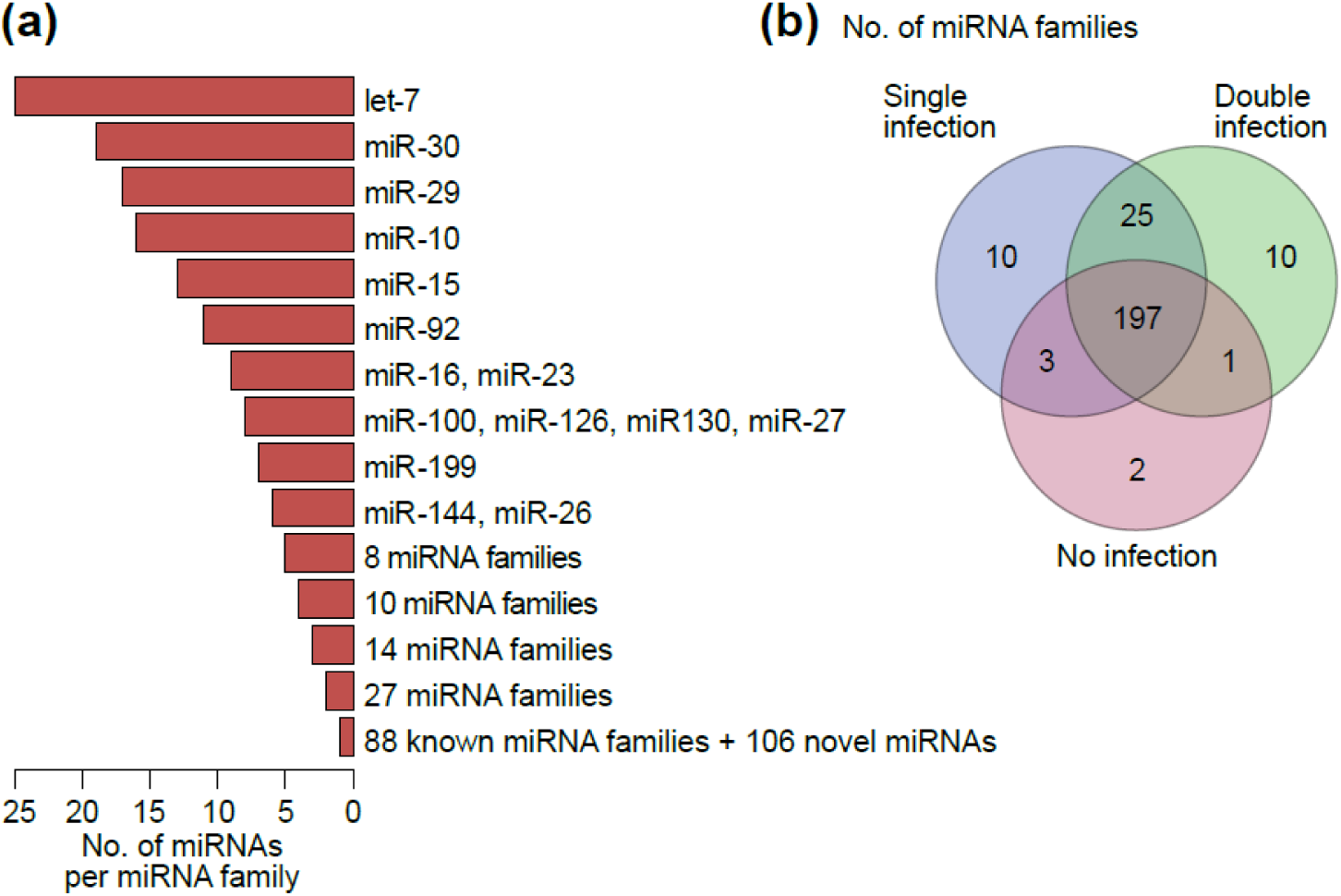
miRNAs families. (**a**) Barplot showing the number of miRNAs in each miRNA family. (**b**) Venn diagram representing the overlap of miRNA families between all three treatment groups.

Out of a total of 248 known miRNA families and potential novel miRNAs, the majority (i.e., 197, 79.4%) were shared across all three treatment groups (Figure 2B). Respectively two (0.8%), ten (4.0%), and ten (4.0%) novel miRNAs were only present in control individuals, single-infected individuals, and double-infected individuals. The two infected groups shared 25 miRNA families that were not present in the control group, totalling 45 (18.1%) miRNA families found only in individuals that were exposed to parasites.

We identified a total of nine DEmirs through pairwise comparisons across all three treatments (Figure 3). Five miRNAs were differentially expressed in single-infected toads compared to uninfected toads, all of them being upregulated (mir-164f, mir-9270, mir-6240, mir-11980, and mir-11987; Figure 3A and D). Four miRNAs were differentially expressed in double-infected toads compared to uninfected toads, all of them being again upregulated (mir-6937-5p, mir-H17, mir-5126, and mir-4154-3p; Figure 3B and E). Out of the nine DEmirs, 5 (i.e., mir-164f, mir-9270, mir-11987, mir-5126, and mir-4154-3p) belonged to miRNA families that were absent (i.e., had <10 cpm) from control individuals. There were no significant differences in miRNA expression between single-infected toads and double-infected toads (Figure 3C). Further, there was no overlap in DEmirs between each pairwise comparison (Figure 3F). None of the 106 potential novel miRNAs were differentially expressed between the three treatment groups.

**Figure 3.**
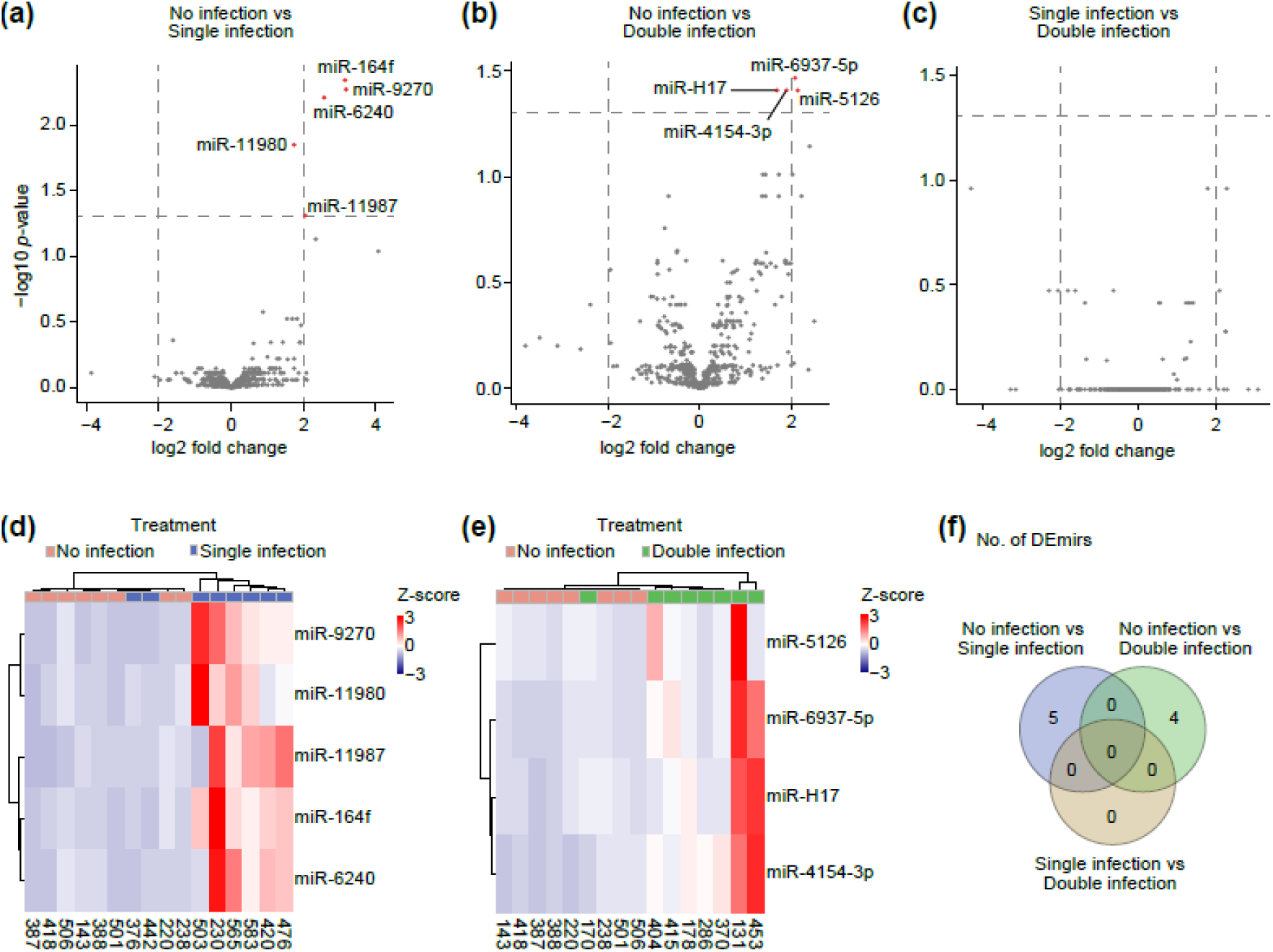
miRNAs differential expression between treatment groups. (**a**–**c**) Volcano plots of DEmirs between (**a**) uninfected and single-infected toads, (**b**) uninfected and double-infected toads, and (**c**) single-infected and double-infected toads. Non-significant miRNAs are represented in grey. (**d**, **e**) Heatmaps of normalised miRNA expression values for all DEmirs between (**d**) uninfected and single-infected toads, and (**e**) uninfected and double-infected toads. Columns correspond to individuals. Rows correspond to miRNAs. Colour depicts Z-score normalised miRNA expression value. (**f**) Venn diagram representing the overlap of DEmirs between all three treatment groups.

### Functional analysis of DEmirs

Prediction analyses revealed 206 ± 195 potential target genes for all DEmirs. GO enrichment analyses of target genes did not reveal any significantly over-represented GO term for all three pairwise comparisons.

Out of nine DEmir families, seven have been linked to biological functions relating to immune responses and/or host-parasite interactions (Table 1). Specifically, mir-11987, mir-164, mir-6240, mir-11980, mir-5126, mir-6937-5p and mir-H17 have been found to be differentially expressed either during pathogenic infections by nematodes, bacteria or viruses, or during cancer or inflammation-related disorders (Abu-Izneid et al., 2021; Escuin et al., 2020; Goher, Hicks, & Liu, 2013; Ma et al., 2016; Ning & Sun, 2020; Palumbo, Faucz, Azevedo, Xekouki, & Iliopoulos, 2013; Wu, Ning, & Sun, 2021; Wu et al., 2020; Yao et al., 2020; Zhao, Hou, & Yan, 2022; Zhu et al., 2018).

**Table 1.** DEmirs involved in immune function and/or host-parasite interactions. U, Uninfected; SI, Single-infected; DI, Double-infected.

| DE-miR | Expression change | Putative role | Reference |
| --- | --- | --- | --- |
| mir-11987 | SI > U<br>Log2FC = 2.05<br>$p = 0.049$ | Upregulated in fish during bacterial and viral infection.<br>Target genes involved in immune response (e.g., *CLN7*, *CCL19*, *GCSF*). | Lewandowski et al., 2022; Lopez-Fabuel et al., 2022; Ning & Sun, 2020; Panopoulos & Watowich, 2008; Wu et al., 2021 |
| mir-6240 | SI > U<br>Log2FC = 2.58<br>$p = 0.006$ | Upregulated in fish during bacterial infection.<br>Upregulated in humans with breast tumours.<br>Target genes involved in immune response (e.g., *ULK1/2*, *CXCL12*). | Cambier et al., 2023; Chan & Tooze, 2009; Escuin et al., 2020; Ning & Sun, 2020 |
| mir-164 | SI > U<br>Log2FC = 3.16<br>$p = 0.005$ | Downregulated in humans with pituitary tumours. | Palumbo et al., 2013 |
| mir-11980 | SI > U<br>Log2FC = 1.75<br>$p = 0.014$ | Upregulated in rats with hypertension. | Wu et al., 2020 |
| mir-6937-5p | DI > U<br>Log2FC = 2.07<br>$p = 0.034$ | Upregulated in mice with retinal dystrophy.<br>Target genes involved in immune response (*ANGPTL1*). | Anasagasti et al., 2020; Gu et al., 2010 |
| mir-5126 | DI > U<br>Log2FC = 2.13<br>$p = 0.039$ | Downregulated in macrophage cells during bacterial infection.<br>Downregulated in mice with inflammation.<br>Target genes involved in drug resistance (e.g., *ABC* and *CYP* genes). | Hollenstein et al., 2007; Ma et al., 2016; Palrasu & Nagini, 2018; Yao et al., 2020; Zhu et al., 2018 |
| mir-H17 | DI > U<br>Log2FC = 1.68<br>$p = 0.039$ | Upregulated in mice and chickens during viral infection. | Du et al., 2015; Goher et al., 2013 |

| DEmir | Expression change | Putative role | Reference |
| --- | --- | --- | --- |
| mir-11987 | SI > U<br>Log2FC = 2.05<br>$p = 0.049$ | Upregulated in fish during bacterial and viral infection.<br>Target genes involved in immune response (e.g., *CLN7*, *CCL19*, *GCSF*). | Lewandowski et al., 2022; Lopez-Fabuel et al., 2022; Ning & Sun, 2020; Panopoulos & Watowich, 2008; Wu et al., 2021 |
| mir-6240 | SI > U<br>Log2FC = 2.58<br>$p = 0.006$ | Upregulated in fish during bacterial infection.<br>Upregulated in humans with breast tumours.<br>Target genes involved in immune response (e.g., *ULK1/2*, *CXCL12*). | Cambier et al., 2023; Chan & Tooze, 2009; Escuin et al., 2020; Ning & Sun, 2020 |
| mir-164 | SI > U<br>Log2FC = 3.16<br>$p = 0.005$ | Downregulated in humans with pituitary tumours. | Palumbo et al., 2013 |
| mir-11980 | SI > U<br>Log2FC = 1.75<br>$p = 0.014$ | Upregulated in rats with hypertension. | Wu et al., 2020 |
| mir-6937-5p | DI > U<br>Log2FC = 2.07<br>$p = 0.034$ | Upregulated in mice with retinal dystrophy.<br>Target genes involved in immune response (*ANGPTL1*). | Anasagasti et al., 2020; Gu et al., 2010 |
| mir-5126 | DI > U<br>Log2FC = 2.13<br>$p = 0.039$ | Downregulated in macrophage cells during bacterial infection.<br>Downregulated in mice with inflammation.<br>Target genes involved in drug resistance (e.g., *ABC* and *CYP* genes). | Hollenstein et al., 2007; Ma et al., 2016; Palrasu & Nagini, 2018; Yao et al., 2020; Zhu et al., 2018 |
| mir-H17 | DI > U<br>Log2FC = 1.68<br>$p = 0.039$ | Upregulated in mice and chickens during viral infection. | Du et al., 2015; Goher et al., 2013 |

### mRNA differential expression

Pairwise comparisons between all three treatment groups revealed only one DEG. *HAUS3* was upregulated in toads from the single infection group compared with toads from the control group (log2 fold change = 4.78; *p*-value = 0.047). No DEG was found between double-infected toads and uninfected toads, nor between single-infected toads and double-infected toads.

To verify that this lack of gene expression differences was not an artefact of removing one toad or of our transcriptomic data, we used the same data to run differential expression analyses according to the toads’ geographic localities in the Australian invasion range (i.e., range-core = Queensland, intermediate = Northern Territory, range-edge = Western Australia). There were large differences in gene expression between range-core toads and toads from both intermediate populations and range-edge populations (respectively 3,458 DEGs and 2,625 DEGs; Figure S3). Toads from intermediate populations and range-edge populations showed a smaller but substantial difference in their spleen transcriptomes (430 DEGs; Figure S3).

## Discussion

Using a co-evolved host-parasite system across an invasive range, we investigated how miRNA and mRNA expression in cane toad hosts responds to infection by their lungworm parasite *R. pseudosphaerocephala*. Our study is the first to characterise miRNAs in cane toads, where we report 540 miRNAs, including over 100 potential novel miRNAs. Given the importance of miRNAs in animal development, physiology and immunity (Carissimi et al., 2009; Carthew, 2006; He & Pan, 2022; O’Brien et al., 2018), and the significance of the cane toad as a model in ecology, evolution and conservation (DeVore, Crossland, Shine, & Ducatez, 2021; Rollins, Richardson, & Shine, 2015; Selechnik et al., 2019; Shine, 2014; Shine et al., 2021), our findings will enable future in-depth investigations of the molecular mechanisms underlying host-parasite co-evolution.

The spleen is a key organ for both innate and adaptive immune responses (Mebius & Kraal, 2005). We showed that miRNA expression in cane toad spleens was upregulated following single or double infection by *R. pseudosphaerocephala* compared to uninfected controls, suggesting that an immune response was mounted. While GO enrichment analyses of DEmirs’ putative target genes did not show any significant over-representation of immunity-related GO terms, seven out of nine DEmir families (i.e., four in uninfected vs single-infected toads, and three in uninfected vs double-infected toads) that were affected have been associated with immune responses and/or host-parasite-related processes. Overall, upregulation of mir-11987, mir-6240, mir-164, mir-11980, mir-6937-5p, mir-5126 and mir-H17 in cane toads infected by *R. pseudosphaerocephala* is consistent with an activation of the toads’ immune system as a direct response to parasite infection.

Increased expression of mir-11987 and mir-6240 may reflect a non-specific immune response from naïve hosts when exposed to lungworm parasites. Both mir-11987 and mir-6240 were upregulated in the spleens of Japanese flounders (*Paralichthys olivaceus*) during an infection by the vibriosis-inducing bacterium *Vibrio anguillarum* (Ning & Sun, 2020). mir-11987 was also upregulated in spleens of *P. olivaceus* during infection by a megalocytivirus (Wu et al., 2021). Further, mir-6240 was upregulated in human primary breast tumours compared to metastases (Escuin et al., 2020). mir-11987 gene targets include *CLN7* (ceroid-lipofuscinosis neuronal protein 7), *CCL19* (C-C motif chemokine 19) and *GCSF* (granulocyte colony-stimulating factor), while mir-6240 targets *ULK1*/*2* (unc-51 like autophagy activating kinase 1/2) and *CXCL12* (C-X-C motif chemokine ligand 12) (Ning & Sun, 2020; Wu et al., 2021). *CLN7* and *ULK1*/*2* regulate anti-bacterial and antiviral innate immune responses through inducing autophagy (Chan & Tooze, 2009; Lopez-Fabuel et al., 2022). *CXCL12* is a homeostatic chemokine involved in cancer and inflammatory diseases (Cambier, Gouwy, & Proost, 2023). *CCL19* is another chemokine playing an important role in the trafficking of immune cells (Lewandowski et al., 2022). Additionally, *GCSF* is a cytokine that controls the production and function of neutrophils, and is thus a key part of the vertebrate immune response (Panopoulos & Watowich, 2008).

Dysregulation of mir-164, mir-11980 and mir-6937-5p has been linked with the pathogenesis of several diseases. mir-164 was downregulated in human pituitary tumours relative to normal tissues, indicating that this dysregulation could be involved in carcinogenesis (Palumbo et al., 2013). mir-11980 was upregulated in brain microvascular pericytes exosomes in rats with hypertension (Wu et al., 2020). Inhibition of mir-6937-5p slowed down visual loss in a mouse model of retinal dystrophy (Anasagasti et al., 2020). mir-6937-5p gene targets include *ANGPTL1* (angiopoietin-1), which is thought to inhibit the activation of macrophage cells (Gu et al., 2010).

Both mir-5126 and mir-H17 have been implicated in host responses to bacterial and viral infections. mir-5126 targets genes involved in drug resistance in the intestinal nematode *Toxocara canis* (Ma et al., 2016), such as *ABC* (ATP-binding cassette transporters) and *CYP* (cytochrome P450) genes (Hollenstein, Dawson, & Locher, 2007; Palrasu & Nagini, 2018). Further, mir-5126 was downregulated in macrophage cells infected by the intracellular bacterial pathogen *Brucella melitensis* (Zhu et al., 2018), and was also downregulated in mice with an intestinal inflammation response (Yao et al., 2020). Finally, mir-H17 was upregulated in viral infections, such as in mice trigeminal ganglia infected by herpes simplex virus (HSV) (Du, Han, Zhou, & Roizman, 2015), or in the spleens of chickens co-infected with two herpesviruses, the pathogenic strain MDV1 and the vaccine strain HVT (Goher et al., 2013).

These results imply that cane toads do not simply tolerate their lungworm parasites. Furthermore, the likely energetic costs of mounting this immune response (Bashir-Tanoli & Tinsley, 2014) confirm that infections by *R. pseudosphaerocephala* have negative fitness effects on the toads, as already suggested by studies reporting detrimental consequences of lungworm infection on cane toad development, immunity and behaviour (Finnerty et al., 2017; Finnerty, Shine, & Brown, 2018; C. Kelehear et al., 2011b; C. Kelehear, Webb, & Shine, 2009b; Selechnik et al., 2017), and showing that cane toads display greater resistance through skin defences when lungworms are larger, live longer and are more infectious (Mayer, Shine, & Brown, 2021; Schlippe Justicia, Mayer, Shine, Shilton, & Brown, 2022).

We found that changes in cane toad miRNA expression following a single infection episode by *R. pseudosphaerocephala* were distinct from those following a double lungworm infection episode. Indeed, there was no overlap among the DEmirs seen in naïve and non-naïve hosts. A number of non-mutually exclusive hypotheses might account for this result. First, this could indicate a mounted immune response from cane toads when facing a previously encountered parasite. Under this scenario, the DEmirs seen in naïve toads would reflect their innate immunity, whereas the DEmirs seen in non-naïve toads would correspond to an adaptive immune response. It has been well established that miRNAs are a key component of adaptive immunity, in particular through regulating the development and function of T and B cells (Carissimi et al., 2009; O’Connell, Rao, Chaudhuri, & Baltimore, 2010). It is thus possible that the specificity of the miRNAs’ transcriptomic profile seen in non-naïve cane toads could be the result of their previous exposure to *R. pseudosphaerocephala* parasites. Indeed, studies have shown in cane toads that short-term and long-term lungworm infections result in different immunological responses (Finnerty et al., 2017). To confirm this mechanistic link, it would be interesting in future studies to closely monitor the expression levels of miRNAs before, during and after repeated exposures to lungworm parasites, together with the expression levels of immune genes in cane toad hosts.

It is also possible that at least some DEmirs seen in infected toads originated from the lungworm parasites themselves. Exchange of miRNAs between parasites and hosts is likely very common (Coakley et al., 2016; Ghalehnoei et al., 2020; He & Pan, 2022), and parasite miRNAs can manipulate host immune cell functions (Liu et al., 2019; Sánchez-López, Trelis, Bernal, & Marcilla, 2021). Schlippe Justicia et al. (2022) have speculated that increased feeding rates in cane toads infected by *R. pseudosphaerocephala*, which is associated with increased growth in the lungworms, could be a sign of host manipulation by this parasite.

Likewise, lungworm infection can alter cane toads’ defecation physiology and behaviour towards warmer and moistier environments, which increases lungworm fertility and the survival of lungworm larvae, and is thus indicative of host manipulation {Finnerty, 2018 #1742}. These effects could be mediated, at least in part, by miRNAs. Although some of our DEmirs have been linked with immune function (see above), it is difficult to ascertain with confidence that these were derived from miRNAs of *R. pseudosphaerocephala*, or that they had immuno-suppressive effects. One promising avenue for future investigations would be determining whether *R. pseudosphaerocephala* release miRNAs-containing exosome-like vesicles, as shown in other parasitic helminths (Wang et al., 2015; Zamanian et al., 2015).

Developments in genomic resources might help identify nematode-specific miRNAs and their gene targets. Third, it is also possible that the non-overlap in DEmirs between both pairwise comparisons has no biological significance and occurred by chance, driven by the low numbers of DEmirs in each pairwise comparison.

In contrast with the miRNAs results, we observed virtually no changes in spleen mRNA expression levels following infection by *R. pseudosphaerocephala*. Indeed, there was only one upregulated gene, i.e., *HAUS3* (HAUS augmin-like complex subunit 3), between single-infected toads and uninfected toads. *HAUS3* regulates centrosome and spindle integrity during cytokinesis and mitosis (Lawo et al., 2009). Its upregulation in toads following infection could be reflective of the cell proliferation phase of the immune response (Medzhitov & Janeway Jr, 2000). More broadly, the mismatch between miRNA and mRNA expression could be due to the relatively short-time window of immune activity following parasite exposure that we would have missed by sampling our tissues two months after lungworm infection, and to the higher stability of miRNAs (Ghalehnoei et al., 2020; Mitchell et al., 2008). Differences in mRNA expression often occur early in a parasitic infection (El-Ashram et al., 2017), especially in the case of a coevolved parasite (Bracamonte, Johnston, Monaghan, & Knopf, 2019). Importantly, these negative results were not an artefact of the data, because analysis of population effects revealed large differences in spleen gene expression between range-core populations and both intermediate and range-edge populations. This effect was of smaller magnitude between intermediate and range-edge populations, a pattern already found in cane toad brain transcriptomes (Yagound et al., 2022).

Factors independent of host-parasite interactions and immune function also may have contributed to the observed differences in miRNA expression between treatment groups. Due to the length of time between the two lungworm exposures for the double-infection group (i.e., 10 months), the developmental stage of the toads at the time of each exposure may have influenced their transcriptomic profiles. It is also possible that the differential number of *R. pseudosphaerocephala* L3 larvae used between the initial and second exposures (i.e., 50–100 vs 30 L3 larvae) affected the response seen in this group compared to the single-infection group. As discussed above, the two-months gap between the final lungworm exposure and tissue collection likely affected the toads’ mRNA expression profiles, and potentially the miRNA response as well.

Helminths are ubiquitous in nature and many parasitic species impact agriculture and human health (Coakley et al., 2016; Maizels, 2020; Sánchez-López et al., 2021). To clarify the mechanisms underlying interactions between these parasites and their hosts, we need genomic studies on the influence of miRNAs on the cross-talk between the organisms involved (Quinzo, Perteguer, Brindley, Loukas, & Sotillo, 2022). We here provide evidence supporting the idea that infection by *R. pseudosphaerocephala* triggers an immune response in their cane toad hosts, and that repeated exposure to lungworm parasites may activate adaptive immune functions. Host manipulation through the release of miRNAs remains an exciting open avenue for future investigations.

## Acknowledgements

Funding for this project was provided by ARC grant DP190100507 awarded to LAR and RS.

## Data Accessibility Statement

The raw RNA-seq data has been made available at the National Center for Biotechnology Information (NCBI) Sequence Read Archive (BioProjects PRJNA949850 and PRJNA952102).

## Author Contributions

TC, BY, LR, GB, RS, and LR designed this project. GB and HE conducted fieldwork. TC and BY analysed data and drafted the manuscript. All authors contributed to editing the final manuscript.

