## Supplementary figures and images for "Infection by the lungworm *Rhabdias pseudosphaerocephala* affects the expression of immune-related microRNAs by its co-evolved host, the cane toad *Rhinella marina*"

### Figure S1

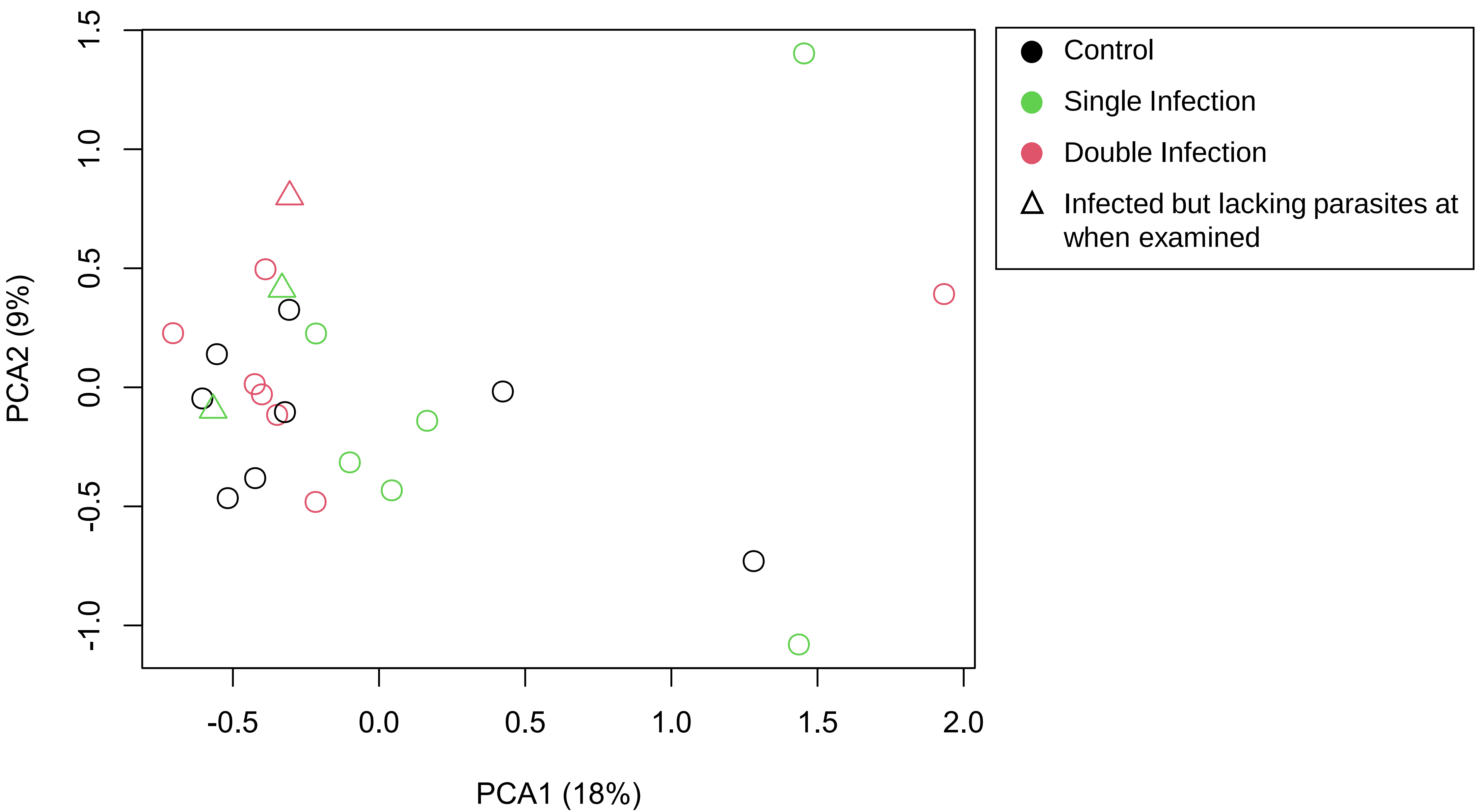

### Figure S2

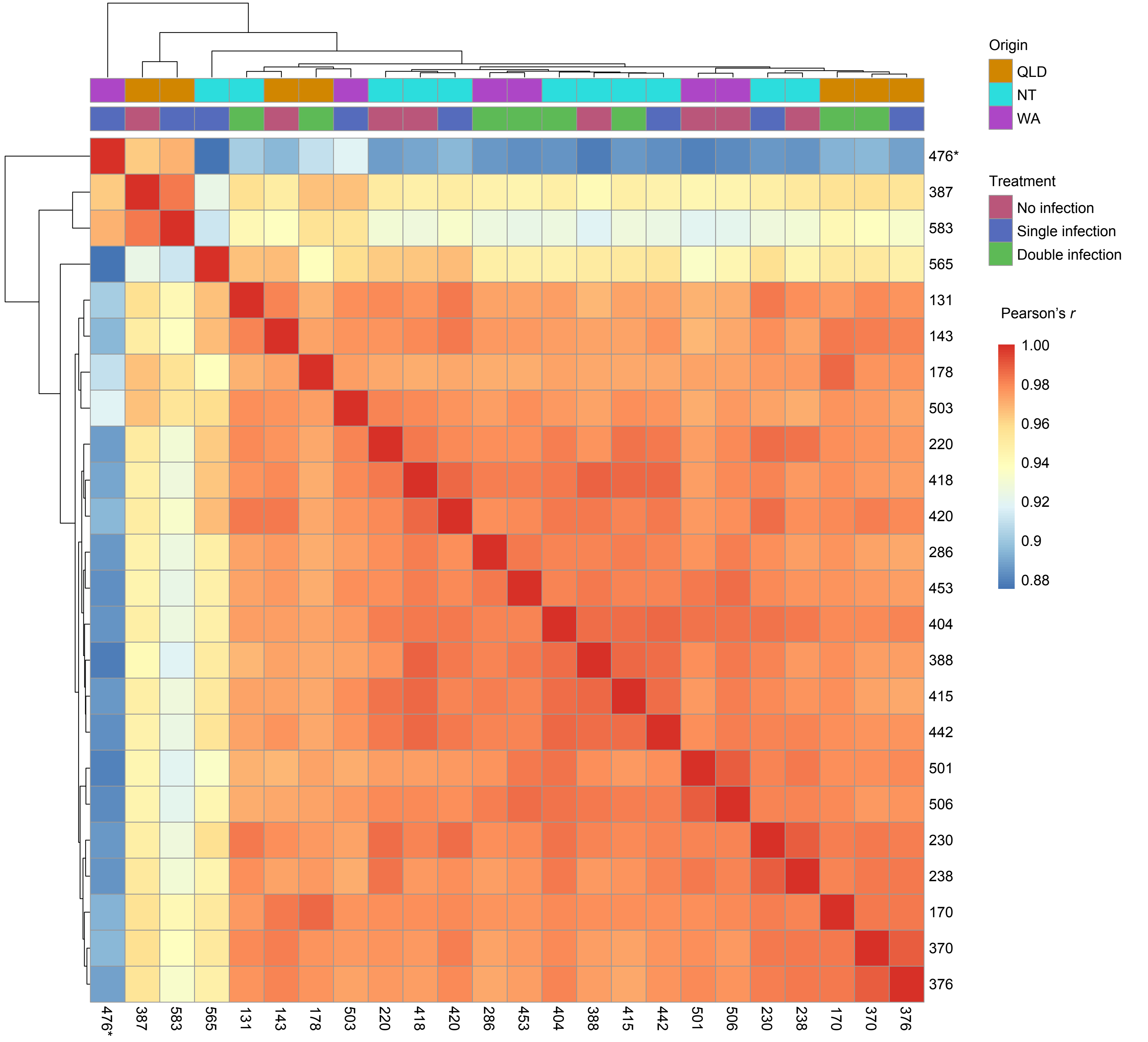

### Figure S3

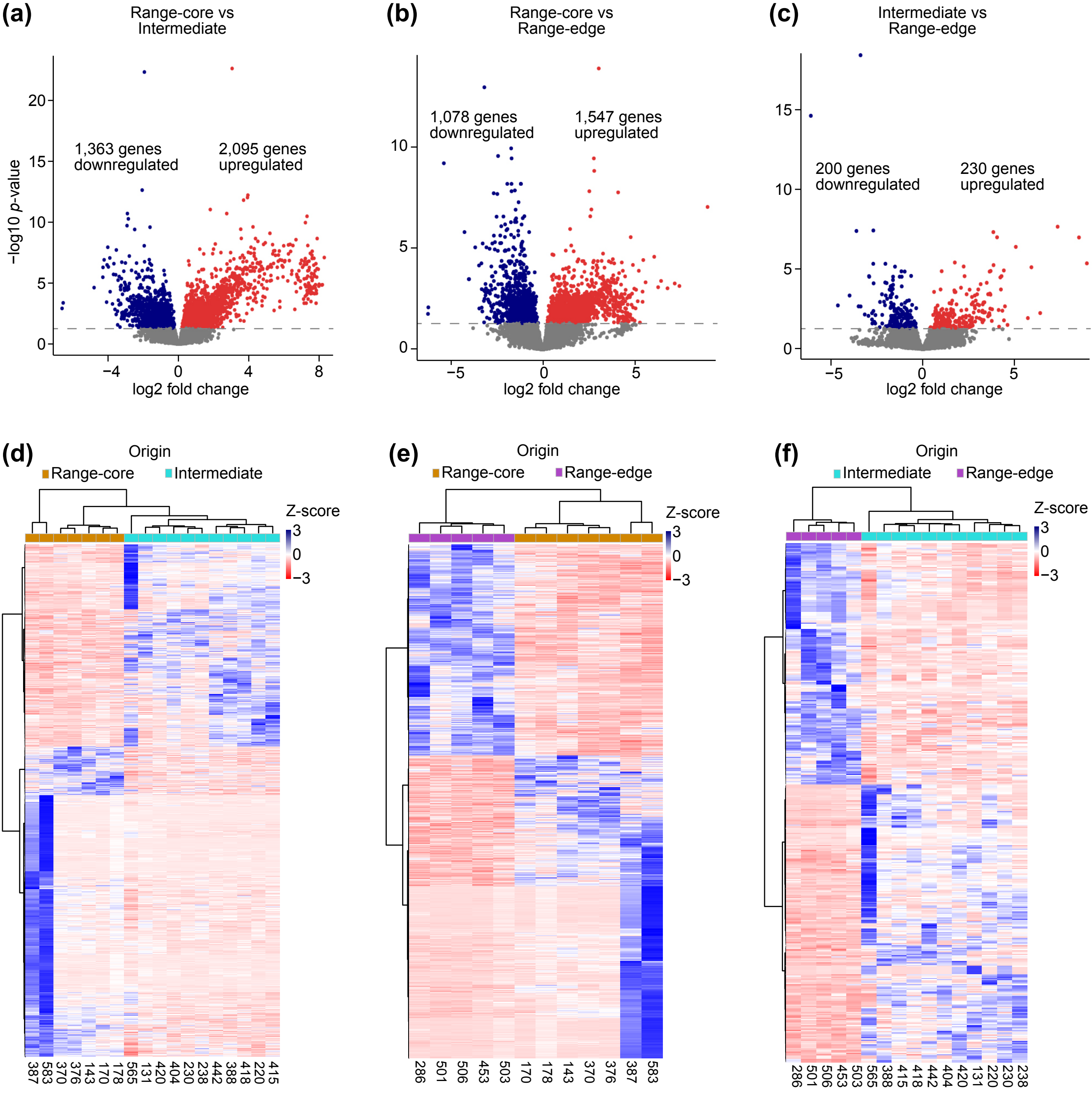
