## Supplementary material for "Infection by the lungworm *Rhabdias pseudosphaerocephala* affects the expression of immune-related microRNAs by its co-evolved host, the cane toad *Rhinella marina*": Table S1

**Table S1.** Samples location and treatment group. QLD, Queensland; NT, Northern Territory; WA, Western Australia; RIN, RNA Integrity Number.

| Individual | Location | Treatment | Worms in lung | Spleen mass (mg) | RIN |
| --- | --- | --- | --- | --- | --- |
| 143 | QLD | Control | 0 | 11 | 9.1 |
| 220 | NT | Control | 0 | 11 | 8.9 |
| 238 | NT | Control | 0 | 7 | 8.0 |
| 387 | QLD | Control | 0 | 14 | 9.0 |
| 388 | NT | Control | 0 | 33 | 9.1 |
| 418 | NT | Control | 0 | 35 | 9.5 |
| 501 | WA | Control | 0 | 1 | 8.2 |
| 506 | WA | Control | 0 | 13 | 9.1 |
| 230 | NT | Single infection | 1 | 8 | 9.0 |
| 376 | QLD | Single infection | 0 | <1 | 8.5 |
| 420 | NT | Single infection | 4 | 9 | 9.2 |
| 442 | NT | Single infection | 7 | 9 | 9.1 |
| 476 | WA | Single infection | 1 | 9 | 9.3 |
| 503 | WA | Single infection | 2 | 1 | 8.3 |
| 565 | NT | Single infection | 1 | 23 | 8.8 |
| 583 | QLD | Single infection | 0 | 9 | 9.2 |
| 131 | NT | Double infection | 1 | 5 | 9.5 |
| 170 | QLD | Double infection | 1 | 15 | 9.1 |
| 178 | QLD | Double infection | 3 | 9 | 9.0 |
| 286 | WA | Double infection | 4 | 11 | 9.4 |
| 370 | QLD | Double infection | 2 | 8 | 8.9 |
| 404 | NT | Double infection | 0 | 20 | 9.6 |
| 415 | NT | Double infection | 4 | 31 | 8.3 |
| 453 | WA | Double infection | 4 | <1 | 8.5 |
