## Supplementary material for "Infection by the lungworm *Rhabdias pseudosphaerocephala* affects the expression of immune-related microRNAs by its co-evolved host, the cane toad *Rhinella marina*": Table S2

**Table S2.** miRNAs sequencing statistics.

| Individual | No. raw reads | No. post-trimming reads (%) | No. mapped reads (%) |
| --- | --- | --- | --- |
| 143 | 16,117,419 | 10,492,261 (65.1) | 9,267,375 (88.3) |
| 220 | 16,408,286 | 11,506,596 (70.1) | 9,865,102 (85.7) |
| 238 | 21,341,273 | 15,529,071 (72.8) | 13,750,582 (88.5) |
| 387 | 18,051,498 | 12,438,051 (68.9) | 10,914,831 (87.8) |
| 388 | 17,700,159 | 11,025,290 (62.3) | 9,751,135 (88.4) |
| 418 | 18,013,896 | 8,560,127 (47.5) | 7,505,040 (87.7) |
| 501 | 13,706,389 | 9,100,109 (66.4) | 8,044,639 (88.4) |
| 506 | 16,399,940 | 9,763,038 (59.5) | 8,484,667 (86.9) |
| 230 | 17,007,272 | 10,880,441 (64.0) | 9,192,597 (84.5) |
| 376 | 16,251,727 | 10,245,536 (63.0) | 9,146,524 (89.3) |
| 420 | 18,639,132 | 12,286,260 (65.9) | 11,025,244 (89.7) |
| 442 | 16,734,826 | 11,169,683 (66.7) | 9,972,560 (89.3) |
| 476 | 19,265,569 | 9,261,448 (48.1) | 7,403,733 (79.9) |
| 503 | 17,468,237 | 4,149,386 (23.8) | 3,413,650 (82.3) |
| 565 | 20,169,899 | 7,285,842 (36.1) | 6,018,803 (82.6) |
| 583 | 17,316,444 | 8,357,053 (48.3) | 6,492,540 (77.7) |
| 131 | 16,771,460 | 4,011,541 (23.9) | 3,361,808 (83.8) |
| 170 | 17,109,077 | 10,615,488 (62.0) | 9,076,566 (85.5) |
| 178 | 16,769,148 | 10,831,501 (64.6) | 9,369,209 (86.5) |
| 286 | 16,558,185 | 11,042,815 (66.7) | 9,590,652 (86.8) |
| 370 | 14,962,276 | 9,059,064 (60.5) | 7,987,797 (88.2) |
| 404 | 17,024,035 | 6,714,993 (39.4) | 5,876,391 (87.5) |
| 415 | 17,688,140 | 6,953,581 (39.3) | 6,120,661 (88.0) |
| 453 | 16,695,598 | 9,794,953 (58.7) | 7,732,332 (78.9) |
