## Supplementary material for "Infection by the lungworm *Rhabdias pseudosphaerocephala* affects the expression of immune-related microRNAs by its co-evolved host, the cane toad *Rhinella marina*": Table S3

**Table S3.** mRNAs sequencing statistics.

| Individual | No. raw reads | No. post-trimming reads (%) | No. mapped reads (%) |
| --- | --- | --- | --- |
| 143 | 44,777,637 | 44,116,414 (98.5) | 26,611,063 (60.3) |
| 220 | 43,912,312 | 43,261,870 (98.5) | 26,401,496 (61.0) |
| 238 | 45,532,712 | 44,837,932 (98.5) | 27,135,808 (60.5) |
| 387 | 39,380,988 | 38,724,596 (98.3) | 24,684,449 (63.7) |
| 388 | 38,161,905 | 37,613,619 (98.6) | 22,615,527 (60.1) |
| 418 | 43,461,467 | 42,751,825 (98.4) | 26,558,281 (62.1) |
| 501 | 48,300,932 | 47,584,965 (98.5) | 28,737,092 (60.4) |
| 506 | 40,042,316 | 39,442,549 (98.5) | 24,400,470 (61.9) |
| 230 | 40,686,692 | 40,023,376 (98.4) | 24,225,701 (60.5) |
| 376 | 45,399,987 | 44,703,756 (98.5) | 26,991,296 (60.4) |
| 420 | 44,441,743 | 43,826,137 (98.6) | 26,901,216 (61.4) |
| 442 | 36,631,356 | 36,102,250 (98.6) | 21,777,487 (60.3) |
| 476 | 48,198,536 | 47,522,847 (98.6) | 33,024,092 (69.5) |
| 503 | 33,534,310 | 33,072,631 (98.6) | 20,491,368 (62.0) |
| 565 | 43,066,996 | 42,413,903 (98.5) | 24,965,462 (58.9) |
| 583 | 38,874,173 | 38,295,544 (98.5) | 24,235,665 (63.3) |
| 131 | 42,214,060 | 41,640,092 (98.6) | 25,369,211 (60.9) |
| 170 | 46,458,455 | 45,753,089 (98.5) | 27,783,108 (60.7) |
| 178 | 41,380,764 | 40,664,198 (98.3) | 24,478,883 (60.2) |
| 286 | 41,102,517 | 40,392,011 (98.3) | 24,217,483 (60.0) |
| 370 | 32,351,447 | 31,901,697 (98.6) | 19,205,573 (60.2) |
| 404 | 37,455,660 | 36,809,359 (98.3) | 22,434,781 (60.9) |
| 415 | 42,745,893 | 42,119,018 (98.5) | 26,101,635 (62.0) |
| 453 | 40,999,580 | 40,397,849 (98.5) | 24,307,568 (60.2) |
