## Supplementary material for "Infection by the lungworm *Rhabdias pseudosphaerocephala* affects the expression of immune-related microRNAs by its co-evolved host, the cane toad *Rhinella marina*": Table 4

**Table S4.** List of all known and potential novel miRNAs identified in this study, and their corresponding miRNA family.

| miRNA | miRNA family |
| --- | --- |
| abu-let-7f | let-7 |
| abu-miR-29c-3p | miR-29 |
| abu-miR-29d | miR-29 |
| aca-miR-139-5p | miR-139 |
| aca-miR-15b-5p | miR-15 |
| aca-miR-215-5p | miR-215 |
| aca-miR-26-2-3p | miR-26 |
| aca-miR-301b-5p | miR-301 |
| aca-miR-9-3-3p | miR-9 |
| aca-miR-99b-5p | miR-99 |
| ami-miR-10c-3p | miR-10 |
| ami-miR-10c-5p | miR-10 |
| ami-miR-1306-3p | miR-1306 |
| ami-miR-497-5p | miR-497 |
| ami-miR-98-5p | miR-98 |
| atr-miR8607 | miR-8607 |
| bbe-miR-29a-3p | miR-29 |
| bbe-miR-375-3p | miR-375 |
| bbe-miR-4868b-3p | miR-4868 |
| bbe-miR-92a-3p | miR-92 |
| bdi-miR164f | miR-164 |
| bfl-miR-92c | miR-92 |
| bta-miR-11980 | miR-11980 |
| bta-miR-11987 | miR-11987 |
| bta-miR-146a | miR-146 |
| bta-miR-150 | miR-150 |
| bta-miR-2340 | miR-2340 |
| bta-miR-2477 | miR-2477 |
| bta-miR-2478 | miR-2478 |
| bta-miR-29d-3p | miR-29 |
| bta-miR-29e | miR-29 |
| bta-miR-30f | miR-30 |
| ccr-miR-10c | miR-10 |
| cgr-miR-1260 | miR-1260 |
| cgr-miR-15a-5p | miR-15 |
| cgr-miR-222-3p | miR-222 |
| chi-let-7b-3p | let-7 |
| chi-miR-16b-5p | miR-16 |
| chi-miR-17-3p | miR-17 |
| chi-miR-29c-3p | miR-29 |
| chi-miR-451-3p | miR-451 |
| cin-miR-126-3p | miR-126 |
| cin-miR-199-3p | miR-199 |
| cin-miR-4154-3p | miR-4154 |
| cin-miR-4171-5p | miR-4171 |
| cin-miR-4185-3p | miR-4185 |
| cin-miR-5606-5p | miR-5606 |
| cli-let-7i-5p | let-7 |
| cli-miR-15b-5p | miR-15 |
| cli-miR-1662-3p | miR-1662 |
| cli-miR-1662-5p | miR-1662 |
| cli-miR-16a-5p | miR-16 |
| cli-miR-194-5p | miR-194 |
| cli-miR-1b-3p | miR-1 |
| cli-miR-23b-3p | miR-23 |
| cli-miR-2970-3p | miR-2970 |
| cli-miR-2970-5p | miR-2970 |
| cli-miR-30c-5p | miR-30 |
| cli-miR-338a-3p | miR-338 |
| cli-miR-449d-5p | miR-449 |
| cli-miR-455-5p | miR-455 |
| cpi-miR-15b-5p | miR-15 |
| cpi-miR-16c-5p | miR-16 |
| cpi-miR-203-3p | miR-203 |
| cpi-miR-454-3p | miR-454 |
| cpi-miR-551-3p | miR-551 |
| cpo-miR-10a-5p | miR-10 |
| cpo-miR-126-5p | miR-126 |
| cpo-miR-15b-5p | miR-15 |
| cpo-miR-30e-3p | miR-30 |
| cpo-miR-425-5p | miR-425 |
| cpo-miR-9-3p | miR-9 |
| ctg1_1 | unknown |
| ctg10297_2 | unknown |
| ctg10326_3 | unknown |
| ctg10355_4 | unknown |
| ctg104_5 | unknown |
| ctg10608_6 | unknown |
| ctg1072_7 | unknown |
| ctg10970_8 | unknown |
| ctg11004_9 | unknown |
| ctg11247_10 | unknown |
| ctg11560_11 | unknown |
| ctg11658_12 | unknown |
| ctg11702_13 | unknown |
| ctg12366_14 | unknown |
| ctg1242_15 | unknown |
| ctg1264_16 | unknown |
| ctg1285_17 | unknown |
| ctg12916_18 | unknown |
| ctg13019_19 | unknown |
| ctg13321_20 | unknown |
| ctg13445_21 | unknown |
| ctg14037_22 | unknown |
| ctg14362_23 | unknown |
| ctg1459_24 | unknown |
| ctg1623_25 | unknown |
| ctg1629_26 | unknown |
| ctg16608_27 | unknown |
| ctg1684_28 | unknown |
| ctg17592_29 | unknown |
| ctg17729_30 | unknown |
| ctg1832_31 | unknown |
| ctg18840_32 | unknown |
| ctg18919_33 | unknown |
| ctg18945_34 | unknown |
| ctg19292_35 | unknown |
| ctg19384_36 | unknown |
| ctg1990_37 | unknown |
| ctg2076_38 | unknown |
| ctg21235_39 | unknown |
| ctg2137_40 | unknown |
| ctg22922_41 | unknown |
| ctg23415_42 | unknown |
| ctg2357_43 | unknown |
| ctg2426_44 | unknown |
| ctg24679_45 | unknown |
| ctg2483_46 | unknown |
| ctg24888_47 | unknown |
| ctg25547_48 | unknown |
| ctg2588_50 | unknown |
| ctg26774_51 | unknown |
| ctg276_52 | unknown |
| ctg27750_53 | unknown |
| ctg28279_54 | unknown |
| ctg28457_55 | unknown |
| ctg2886_56 | unknown |
| ctg2897_57 | unknown |
| ctg3096_58 | unknown |
| ctg3108_59 | unknown |
| ctg3117_60 | unknown |
| ctg314_61 | unknown |
| ctg3355_62 | unknown |
| ctg3381_63 | unknown |
| ctg3427_64 | unknown |
| ctg3461_65 | unknown |
| ctg3519_66 | unknown |
| ctg3614_67 | unknown |
| ctg378_68 | unknown |
| ctg3843_68 | unknown |
| ctg3979_69 | unknown |
| ctg4047_70 | unknown |
| ctg4309_71 | unknown |
| ctg4334_72 | unknown |
| ctg4450_73 | unknown |
| ctg5051_74 | unknown |
| ctg5067_75 | unknown |
| ctg5267_76 | unknown |
| ctg541_77 | unknown |
| ctg5416_78 | unknown |
| ctg5421_79 | unknown |
| ctg5577_80 | unknown |
| ctg5586_81 | unknown |
| ctg569_82 | unknown |
| ctg5841_83 | unknown |
| ctg5957_84 | unknown |
| ctg5986_85 | unknown |
| ctg6266_86 | unknown |
| ctg6283_87 | unknown |
| ctg634_88 | unknown |
| ctg642_89 | unknown |
| ctg6585_90 | unknown |
| ctg6742_91 | unknown |
| ctg6905_92 | unknown |
| ctg6977_93 | unknown |
| ctg7106_94 | unknown |
| ctg7146_95 | unknown |
| ctg7251_96 | unknown |
| ctg7349_97 | unknown |
| ctg76_98 | unknown |
| ctg786_99 | unknown |
| ctg8013_01 | unknown |
| ctg8737_02 | unknown |
| ctg9093_03 | unknown |
| ctg923_04 | unknown |
| ctg9263_05 | unknown |
| ctg9463_06 | unknown |
| ctg966_07 | unknown |
| dma-miR-10a | miR-10 |
| dme-miR-4969-5p | miR-4969 |
| dno-miR-215-3p | miR-215 |
| dno-miR-23a-3p | miR-23 |
| dno-miR-24b-3p | miR-24 |
| dno-miR-410-5p | miR-410 |
| dqu-miR-92a-3p | miR-92 |
| dre-let-7g | let-7 |
| dre-miR-101a | miR-101 |
| dre-miR-107b | miR-107 |
| dre-miR-126b-3p | miR-126 |
| dre-miR-16c-5p | miR-16 |
| dre-miR-199-3-3p | miR-199 |
| dre-miR-202-5p | miR-202 |
| dre-miR-27a-3p | miR-27 |
| dre-miR-27b-3p | miR-27 |
| dre-miR-338-3p | miR-338 |
| dre-miR-455-2-5p | miR-455 |
| dre-miR-456 | miR-456 |
| dre-miR-457b-5p | miR-457 |
| dvi-miR-10-5p | miR-10 |
| eca-miR-130b | miR-130 |
| eca-miR-8975 | miR-8975 |
| eel-miR-11055-5p | miR-11055 |
| efu-let-7c | let-7 |
| efu-let-7d | let-7 |
| efu-let-7e | let-7 |
| efu-let-7f | let-7 |
| efu-miR-101 | miR-101 |
| efu-miR-103b | miR-103 |
| efu-miR-107 | miR-107 |
| efu-miR-125b | miR-125 |
| efu-miR-126 | miR-126 |
| efu-miR-128a | miR-128 |
| efu-miR-128b | miR-128 |
| efu-miR-130 | miR-130 |
| efu-miR-133-3p | miR-133 |
| efu-miR-138a | miR-138 |
| efu-miR-138b | miR-138 |
| efu-miR-143 | miR-143 |
| efu-miR-145 | miR-145 |
| efu-miR-155 | miR-155 |
| efu-miR-16 | miR-16 |
| efu-miR-17 | miR-17 |
| efu-miR-18 | miR-18 |
| efu-miR-181e | miR-181 |
| efu-miR-19 | miR-19 |
| efu-miR-192 | miR-192 |
| efu-miR-196 | miR-196 |
| efu-miR-199 | miR-199 |
| efu-miR-20 | miR-20 |
| efu-miR-200a | miR-200 |
| efu-miR-206 | miR-206 |
| efu-miR-214 | miR-214 |
| efu-miR-218a | miR-218 |
| efu-miR-218b | miR-218 |
| efu-miR-22 | miR-22 |
| efu-miR-223 | miR-223 |
| efu-miR-23a | miR-23 |
| efu-miR-23b | miR-23 |
| efu-miR-25 | miR-25 |
| efu-miR-26a | miR-26 |
| efu-miR-26c | miR-26 |
| efu-miR-27b | miR-27 |
| efu-miR-29a | miR-29 |
| efu-miR-29b | miR-29 |
| efu-miR-29c | miR-29 |
| efu-miR-30a | miR-30 |
| efu-miR-30b | miR-30 |
| efu-miR-30e | miR-30 |
| efu-miR-339 | miR-339 |
| efu-miR-34a | miR-34 |
| efu-miR-383 | miR-383 |
| efu-miR-455 | miR-455 |
| efu-miR-499 | miR-499 |
| efu-miR-7b | miR-7 |
| efu-miR-9270 | miR-9270 |
| efu-miR-92a | miR-92 |
| efu-miR-93 | miR-93 |
| efu-miR-99a | miR-99 |
| egr-miR-10240-5p | miR-10240 |
| fru-miR-210 | miR-210 |
| gga-miR-126-3p | miR-126 |
| gga-miR-1306-5p | miR-1306 |
| gga-miR-142-5p | miR-142 |
| gga-miR-143-3p | miR-143 |
| gga-miR-144-3p | miR-144 |
| gga-miR-145-5p | miR-145 |
| gga-miR-146c-5p | miR-146 |
| gga-miR-1599 | miR-1599 |
| gga-miR-1692 | miR-1692 |
| gga-miR-18b-3p | miR-18 |
| gga-miR-21-3p | miR-21 |
| gga-miR-460b-3p | miR-460 |
| gga-miR-460b-5p | miR-460 |
| gga-miR-6670-5p | miR-6670 |
| ggo-miR-454 | miR-454 |
| gmo-let-7g-3p | let-7 |
| gmo-miR-100b-5p | miR-100 |
| gmo-miR-101a-3p | miR-101 |
| gmo-miR-11202-5p | miR-11202 |
| gmo-miR-126-3p | miR-126 |
| gmo-miR-130b-3p | miR-130 |
| gmo-miR-192-5p | miR-192 |
| gmo-miR-2184-5p | miR-2184 |
| gmo-miR-221-5p | miR-221 |
| gmo-miR-223a-5p | miR-223 |
| gmo-miR-29b-2-5p | miR-29 |
| gmo-miR-730-5p | miR-730 |
| gmo-miR-7550-5p | miR-7550 |
| hco-miR-5991 | miR-5991 |
| hhi-let-7j | let-7 |
| hhi-miR-183 | miR-183 |
| hhi-miR-301 | miR-301 |
| hme-let-7 | let-7 |
| hpo-miR-100-5p | miR-100 |
| hsa-miR-10396b-3p | miR-10396 |
| hsa-miR-10400-5p | miR-10400 |
| hsa-miR-12135 | miR-12135 |
| hsa-miR-22-5p | miR-22 |
| hsa-miR-221-3p | miR-221 |
| hsa-miR-3182 | miR-3182 |
| hsa-miR-4454 | miR-4454 |
| hsa-miR-4492 | miR-4492 |
| hsa-miR-7-5p | miR-7 |
| hsa-miR-7704 | miR-7704 |
| hsa-miR-7847-3p | miR-7847 |
| hsa-miR-7977 | miR-7977 |
| hsv1-miR-H17 | miR-H17 |
| ipu-miR-24 | miR-24 |
| ipu-miR-26b | miR-26 |
| ipu-miR-27d | miR-27 |
| ipu-miR-99b | miR-99 |
| lva-let-7-5p | let-7 |
| mdo-miR-10b-5p | miR-10 |
| mdo-miR-122-5p | miR-122 |
| mdo-miR-144-3p | miR-144 |
| mdo-miR-150-5p | miR-150 |
| mdo-miR-15a-5p | miR-15 |
| mdo-miR-22-3p | miR-22 |
| mdo-miR-460-3p | miR-460 |
| mdo-miR-7384-5p | miR-7384 |
| mml-miR-145-3p | miR-145 |
| mml-miR-20a-3p | miR-20 |
| mml-miR-363-5p | miR-363 |
| mmr-miR-148b | miR-148 |
| mmr-miR-29c | miR-29 |
| mmr-miR-30a | miR-30 |
| mmu-let-7j | let-7 |
| mmu-miR-1187 | miR-1187 |
| mmu-miR-12188-5p | miR-12188 |
| mmu-miR-126b-5p | miR-126 |
| mmu-miR-145a-3p | miR-145 |
| mmu-miR-2137 | miR-2137 |
| mmu-miR-30a-3p | miR-30 |
| mmu-miR-3964 | miR-3964 |
| mmu-miR-466i-5p | miR-466 |
| mmu-miR-5126 | miR-5126 |
| mmu-miR-6239 | miR-6239 |
| mmu-miR-6240 | miR-6240 |
| mmu-miR-6937-5p | miR-6937 |
| mmu-miR-93-5p | miR-93 |
| mse-miR-2779 | miR-2779 |
| nle-miR-30b | miR-30 |
| nve-miR-100-5p | miR-100 |
| oan-miR-1357 | miR-1357 |
| oan-miR-1386 | miR-1386 |
| oan-miR-1417-3p | miR-1417 |
| oan-miR-190b | miR-190 |
| oan-miR-215-3p | miR-215 |
| oan-miR-223-3p | miR-223 |
| oan-miR-23b-3p | miR-23 |
| oan-miR-31-5p | miR-31 |
| oan-miR-451 | miR-451 |
| oan-miR-92b | miR-92 |
| ocu-let-7b-3p | let-7 |
| ocu-miR-10b-3p | miR-10 |
| ocu-miR-10b-5p | miR-10 |
| ocu-miR-130a-3p | miR-130 |
| ocu-miR-137-3p | miR-137 |
| ocu-miR-16a-5p | miR-16 |
| ocu-miR-18b-5p | miR-18 |
| ocu-miR-20b-5p | miR-20 |
| ocu-miR-21-5p | miR-21 |
| ocu-miR-215-3p | miR-215 |
| oga-miR-23b | miR-23 |
| oga-miR-98 | miR-98 |
| oha-miR-100-5p | miR-100 |
| oha-miR-10b-5p | miR-10 |
| oha-miR-10c-5p | miR-10 |
| oha-miR-122-5p | miR-122 |
| oha-miR-129b-5p | miR-129 |
| oha-miR-181a-5p | miR-181 |
| oha-miR-1a-3p | miR-1 |
| oha-miR-21-3p | miR-21 |
| oha-miR-27b-5p | miR-27 |
| oha-miR-30d-3p | miR-30 |
| oha-miR-363-3p | miR-363 |
| oha-miR-429-3p | miR-429 |
| ola-miR-142 | miR-142 |
| oni-miR-10792 | miR-10792 |
| oni-miR-10926 | miR-10926 |
| oni-miR-10955 | miR-10955 |
| oni-miR-10965 | miR-10965 |
| osa-miR5072 | miR-5072 |
| pab-miR11504 | miR-11504 |
| pal-miR-142-3p | miR-142 |
| pal-miR-181a-5p | miR-181 |
| pal-miR-194b-5p | miR-194 |
| pal-miR-30d-5p | miR-30 |
| pal-miR-9226-5p | miR-9226 |
| pal-miR-9993a-3p | miR-9993 |
| pal-miR-9993b-3p | miR-9993 |
| pal-miR-9995-3p | miR-9995 |
| pbv-miR-106-5p | miR-106 |
| pbv-miR-125a-5p | miR-125 |
| pbv-miR-130a-3p | miR-130 |
| pbv-miR-130c-3p | miR-130 |
| pbv-miR-1388-3p | miR-1388 |
| pbv-miR-143-5p | miR-143 |
| pbv-miR-155a-5p | miR-155 |
| pbv-miR-15a-5p | miR-15 |
| pbv-miR-16a-5p | miR-16 |
| pbv-miR-199-5p | miR-199 |
| pbv-miR-204-5p | miR-204 |
| pbv-miR-30d-3p | miR-30 |
| pbv-miR-99a-5p | miR-99 |
| pca-miR-971-5p | miR-971 |
| peu-miR2916 | miR-2916 |
| pha-miR-30e | miR-30 |
| pma-let-7c | let-7 |
| pma-let-7d | let-7 |
| pma-miR-126 | miR-126 |
| pma-miR-143-3p | miR-143 |
| pma-miR-15a | miR-15 |
| pma-miR-16-5p | miR-16 |
| pma-miR-192-5p | miR-192 |
| pma-miR-194-5p | miR-194 |
| pma-miR-20a-5p | miR-20 |
| pma-miR-210 | miR-210 |
| pma-miR-23a-3p | miR-23 |
| pma-miR-29a-3p | miR-29 |
| pma-miR-29b | miR-29 |
| pma-miR-29d-3p | miR-29 |
| pma-miR-30b | miR-30 |
| pma-miR-30c-5p | miR-30 |
| pma-miR-30d | miR-30 |
| pma-miR-4600 | miR-4600 |
| pma-miR-551 | miR-551 |
| pmi-miR-133-3p | miR-133 |
| pmi-miR-33-5p | miR-33 |
| pmi-miR-92c-3p | miR-92 |
| pny-miR-106 | miR-106 |
| pny-miR-130b-3p | miR-130 |
| pny-miR-140 | miR-140 |
| pny-miR-7147 | miR-7147 |
| pol-miR-144-3p | miR-144 |
| ppt-miR894 | miR-894 |
| ppy-miR-144 | miR-144 |
| ppy-miR-199b-3p | miR-199 |
| prd-let-7-5p | let-7 |
| prd-miR-100-5p | miR-100 |
| pte-miR-100a-5p | miR-100 |
| pte-miR-125a-5p | miR-125 |
| pte-miR-210-3p | miR-210 |
| pte-miR-92c-3p | miR-92 |
| ptr-miR-1260b | miR-1260 |
| ptr-miR-143 | miR-143 |
| rno-miR-155-5p | miR-155 |
| rno-miR-17-1-3p | miR-17 |
| rno-miR-215 | miR-215 |
| rno-miR-551b-3p | miR-551 |
| rno-miR-99a-3p | miR-99 |
| sbo-let-7b | let-7 |
| sbo-miR-191 | miR-191 |
| sko-miR-92a | miR-92 |
| spu-miR-10 | miR-10 |
| spu-miR-92a | miR-92 |
| spu-miR-92b-3p | miR-92 |
| ssa-let-7f-5p | let-7 |
| ssa-let-7j-5p | let-7 |
| ssa-miR-10d-5p | miR-10 |
| ssa-miR-142b-3p | miR-142 |
| ssa-miR-144-5p | miR-144 |
| ssa-miR-15a-5p | miR-15 |
| ssa-miR-192a-2-3p | miR-192 |
| ssa-miR-205a-2-3p | miR-205 |
| ssa-miR-214-3p | miR-214 |
| ssa-miR-21a-5p | miR-21 |
| ssa-miR-23a-5p | miR-23 |
| ssa-miR-23b-3p | miR-23 |
| ssa-miR-26d-5p | miR-26 |
| ssa-miR-29b-1-5p | miR-29 |
| ssa-miR-30d-5p | miR-30 |
| ssa-miR-338a-5p | miR-338 |
| ssa-miR-499a-5p | miR-499 |
| ssc-miR-122-5p | miR-122 |
| tca-miR-100-5p | miR-100 |
| tch-let-7a-5p | let-7 |
| tch-miR-26a-2-5p | miR-26 |
| tch-miR-27a-3p | miR-27 |
| tgu-miR-2970-3p | miR-2970 |
| tni-miR-142b | miR-142 |
| tni-miR-15b | miR-15 |
| xla-let-7a-3p | let-7 |
| xla-let-7b-3p | let-7 |
| xla-miR-100-2-3p | miR-100 |
| xla-miR-103b-3p | miR-103 |
| xla-miR-10a-3p | miR-10 |
| xla-miR-10a-5p | miR-10 |
| xla-miR-130a-3p | miR-130 |
| xla-miR-135-5p | miR-135 |
| xla-miR-140-3p | miR-140 |
| xla-miR-140-5p | miR-140 |
| xla-miR-144-3p | miR-144 |
| xla-miR-148a-3p | miR-148 |
| xla-miR-15b-3p | miR-15 |
| xla-miR-15b-5p | miR-15 |
| xla-miR-15c-3p | miR-15 |
| xla-miR-17-3p | miR-17 |
| xla-miR-17-5p | miR-17 |
| xla-miR-192-3p | miR-192 |
| xla-miR-193-5p | miR-193 |
| xla-miR-194-3p | miR-194 |
| xla-miR-199-3p | miR-199 |
| xla-miR-19a-3p | miR-19 |
| xla-miR-19b-3p | miR-19 |
| xla-miR-19b-5p | miR-19 |
| xla-miR-210-5p | miR-210 |
| xla-miR-22-3p | miR-22 |
| xla-miR-22-5p | miR-22 |
| xla-miR-25-5p | miR-25 |
| xla-miR-27b-5p | miR-27 |
| xla-miR-29c-5p | miR-29 |
| xla-miR-30c-3p | miR-30 |
| xla-miR-30e-5p | miR-30 |
| xla-miR-338-5p | miR-338 |
| xla-miR-33b-3p | miR-33 |
| xla-miR-33b-5p | miR-33 |
| xla-miR-375-3p | miR-375 |
| xla-miR-425-2-3p | miR-425 |
| xla-miR-425-3p | miR-425 |
| xla-miR-499-3p | miR-499 |
| xla-miR-92a-4-5p | miR-92 |
| xtr-let-7b | let-7 |
| xtr-let-7e | let-7 |
| xtr-miR-106 | miR-106 |
| xtr-miR-10c | miR-10 |
| xtr-miR-16b | miR-16 |
| xtr-miR-181a-1-3p | miR-181 |
| xtr-miR-181a-2-3p | miR-181 |
| xtr-miR-193 | miR-193 |
| xtr-miR-199b | miR-199 |
| xtr-miR-2184 | miR-2184 |
| xtr-miR-2188 | miR-2188 |
| xtr-miR-24b | miR-24 |
| xtr-miR-27c | miR-27 |
| xtr-miR-29d | miR-29 |
| xtr-miR-460a-3p | miR-460 |
| xtr-miR-499 | miR-499 |
| xtr-miR-9b-3p | miR-9 |
| xtr-miR-9b-5p | miR-9 |
